# Robust Induction of Spontaneous Recurrent Seizures and Hippocampal Sclerosis with Synthetic Kainic Acid in the Intrahippocampal Kainate Model

**DOI:** 10.64898/2026.09.24.754069

**Authors:** Anjali R. Grillo, Paige O’Gorman, Monique Krummradt, Marikate Kenny, Kevin J. Staley, Kyle P. Lillis, Lauren A. Lau

## Abstract

Intrahippocampal kainate (IHK) is a well-established mouse model of temporal lobe epilepsy, in which focal hippocampal injection of kainic acid induces excitotoxic injury followed by the development of chronic spontaneous seizures. Commercial sources of kainic acid (KA) have increasingly transitioned from naturally derived to synthetic KA, which has introduced a potential source of variability to the implementation of the IHK model. Here, we evaluated injection parameters for synthetic KA, with the goal of identifying conditions that reliably produce spontaneous seizures while minimizing mortality. We found that intrahippocampal injection of 200nL of 5mM synthetic KA resulted in low mortality compared to 50-100nL of 20mM synthetic KA (a dosing range commonly used with naturally derived KA), while robustly inducing TLE-like hippocampal sclerosis and recurrent spontaneous electroclinical seizures. We further demonstrated high inter-animal variability in both seizure frequency and temporal clustering of seizures across animals receiving either injection protocol (200nL of 5mM or 50nL of 20mM). Together, these findings provide a framework for implementing the IHK model with synthetic KA and support its continued use to advance our understanding of the cellular and network mechanisms underlying temporal lobe epilepsy.

## Introduction

Epilepsy is estimated to affect 1.1% of the adult population^1^, with temporal lobe epilepsy (TLE) accounting for ∼20% of epilepsy cases^2^. TLE is associated with spontaneous focal onset seizures and often with hippocampal sclerosis. TLE has a high rate of pharmacoresistance, with estimates as high as 50-75% of patients experiencing poor seizure control with antiseizure medications alone, particularly among those with hippocampal sclerosis^3-5^. Addressing this major unmet clinical need will require investigations using animal models that recapitulate the key pathophysiological features of TLE. Typically, rodent models of TLE induce brain injury via administration of a chemoconvulsant to provoke a period of status epilepticus. This insult is then followed by a latent period before the emergence of spontaneous, recurrent seizures. Kainic acid (KA), a potent glutamate receptor agonist^6^, can be administered either systemically or directly into the brain to trigger status epilepticus and epileptogenesis^7^. The intrahippocampal kainate (IHK) mouse model, in which a small volume of KA is administered directly to the hippocampus, has been widely adopted in studies of TLE due to the focal hippocampal injury and robust development of spontaneous recurrent seizures that resemble those seen in human TLE^8-14^.

However, considerable variability exists across IHK induction protocols and in the resulting outcomes, including severity of status epilepticus, mortality, spontaneous seizure frequency and hippocampal histopathology^8-14^. Such variability can complicate comparisons across studies and may limit the reproducibility and interpretability of experimental findings. Establishing standardized and reproducible approaches to IHK induction will maximize the utility of this model for mechanistic and therapeutic studies. One source of variability in the IHK model is the form of kainic acid used in the intrahippocampal injection. Kainic acid (KA) can be obtained either by extraction and purification from naturally occurring sources or through chemical synthesis. Historically, kainic acid was obtained by extraction from the *Digenea simplex* seaweed^15^. Prior to 1995, the sole commercial provider of kainic acid was a Taiwanese company, King Tom Pharmaceutical. When the company discontinued production, a temporary shortage emerged in the supply of kainic acid^16^. In response, Tocris Bioscience (Bristol, UK) began sourcing *Digenea simplex* for the extraction and purification of kainic acid and became one of the major suppliers of natural KA used in the IHK model^8-11^. Recent advances in the chemical synthesis of kainic acid have improved the yield and reduced the total number of synthetic steps^17^, making chemical synthesis a more viable source of commercially available KA. In 2024, Tocris discontinued naturally derived kainic acid and transitioned to exclusively supplying synthetic kainic acid. In this study, we have performed a systematic optimization of synthetic kainic acid for use in the IHK mouse model. This work was motivated by our observation that intrahippocampal injection of synthetic kainic acid resulted in a steep increase in acute model mortality when administered using protocols developed for natural KA (50-100nL of 20mM KA). We propose that synthetic KA should be delivered in a more diluted format (200nL of 5mM) to minimize mortality, while still preserving the development of hippocampal sclerosis and spontaneous recurrent seizures.

## Methods

### Animals

All animal protocols were approved by the Massachusetts General Hospital’s Institutional Animal Care and Use Committee (IACUC). Adult mice (>8-weeks old) of both sexes were used in this study. Two genotypes were included, wild-type C57BL6/J and DLX/tdtomato mice (generated by crossing DLX5/6-cre mice on a mixed C57/CD1 background with Ai14(RCL-tdT) animals on a C57BL6/J background). The study was not designed or powered to assess sex-or genotype-dependent differences in IHK outcomes. However, groups were balanced as feasible for these factors to minimize confounding comparisons between experimental conditions. Sex-and genotype-dependent effects will be investigated in future studies.

### Kainic Acid Preparation

Synthetic kainic acid (KA) was obtained from Tocris Bioscience (#7065). KA solutions were prepared fresh on the day of IHK surgery in 0.9% sterile saline and sonicated for 30 min. KA solutions were discarded at the end of the day.

### Intrahippocampal Injection of KA

Briefly, animals were anesthetized with isoflurane, at 4-5% for induction and 1.5-3% for maintenance. Eye ointment was applied and animals were kept warm on a heating pad during the procedure. The scalp was shaved and cleaned with Betadine and 70% ethanol. A midline incision was made to expose the skull surface. A burr hole was drilled using a 0.9mm drill bit above the right hippocampus. Stereotaxic coordinates for the burr hole were -2.5 mm anterior-posterior (A-P) to Bregma and 1.5 mm medio-lateral (M-L). A 0.5µL Neuros Hamilton Syringe was lowered 1.5 mm from the brain surface to the right dorsal hippocampus (−1.5 D-V). The injection was performed with a programable syringe pump (Harvard Apparatus) at a rate of 25nL/min. At the end of the injection period, the needle was slowly removed. The incision was then closed with sutures.

#### Status Epilepticus Monitoring

Animals were continuously monitored during the period of KA-induced status epilepticus (SE), and the severity of behavioral seizures were scored based on a modified Racine scale^18,19^, as noted in Table 1. In accordance with humane endpoints developed with guidance from IACUC, animals that experienced a continuous Stage 5+ seizure lasting >30 minutes or a Stage 6 seizure lasting >10 minutes were euthanized. Animals were monitored for a minimum of 3 hours post-KA injection. Monitoring ended when normal behavior resumed (defined by exploration and/or grooming) with at least 30 minutes since the last observed seizure. Animals that did not have at least one Stage 5+ seizure during SE were excluded from the study.

**Table 1:** Behavioral criteria for seizure severity staging

| Stage | Description |
| --- | --- |
| 1 | frozen stance and/or mastication/facial movements |
| 2 | Head nodding |
| 3 | Unilateral forelimb clonus |
| 4 | Bilateral forelimb clonus with rearing |
| 5 | Rearing and falling |
| 6 | Wild running and/or jumping |

#### 72-hour Monitoring

Animals were checked twice a day for 72 hours post-KA injection. In accordance with humane endpoints developed with guidance from IACUC, animals that lost >20% of pre-surgical body weight were euthanized. Excessive weight loss was presumed to be associated with ongoing seizure activity as no sham intrahippocampal saline injected animals required euthanasia.

### Electroencephalogram (EEG) implants and recordings

At least one-week post-IHK injection, a subset of animals underwent implantation for 2-EEG/1-EMG recordings, with prefabricated headmounts (Pinnacle Technology, 8201-SS). Briefly, animals were anesthetized with isoflurane, at 4-5% for induction and 1.5-3% for maintenance. The scalp was shaved and cleaned with Betadine and 70% ethanol. A midline incision was made to expose the skull surface. The skull surface was cleaned and dried. Four burr holes were drilled for the placement of stainless-steel screws serving as the EEG electrodes. The two active EEG electrodes were placed bilaterally over the posterior cerebral cortex. Reference and ground electrode were positioned over the frontal cortex. EMG wires were placed in the nuchal muscles. Screws were connected to the headmount with a silver epoxy and the headmount was fixed into place using dental cement. The incision was then closed with sutures. Animals were given at least one week to recover before EEG recordings began.

Animals were transferred to specialty housing to accommodate free movement during tethered recordings, under standard light conditions, with ad libitum access to food and water. Individual animals underwent continuous video-EEG recordings that lasted 7-9 days. Chronic recordings were acquired using at 1kHz using a 10x gain preamplifier attached to the data conditioning and acquisition system (Pinnacle Technology) with simultaneous video recordings. Data was recorded with the Pinnacle Sirenia software and analyzed with Sirenia Seizure Pro.

### Seizure Detection

Putative seizure events were initially identified via automated analysis of the EEG signal based on 1-sec line-length measurements, in which epochs that exceeded the mean + 3 standard deviations threshold were selected for visual review by an investigator. (A subset of data was visually screening in its entirety to ensure that line-length based thresholding did not miss any seizures). Events were classified as seizures if they were at least 10 seconds in duration and exhibited a sustained increase in EEG amplitude accompanied by characteristic progression in waveform morphology and frequency over the course of the event^8,20^.

### Immunohistochemistry

#### Tissue Preparation

Animals were deeply anesthetized with isoflurane and then transcardially perfused with phosphate buffered saline (PBS), followed by 4% paraformaldehyde (PFA). Brains were extracted and post-fixed overnight in 4% PFA at 4 °C then cryoprotected sequentially in 15% and 30% sucrose in PBS at 4 °C until fully submerged.

Cryoprotected brains were embedded in OCT compound and oriented coronally (rostrocaudal axis, olfactory bulbs to cerebellum) on a cryostat chuck. Ipsilateral (IL, side of KA injection) and contralateral (CL) hippocampus were distinguished by a small hole marked on the bottom of the CL side prior to sectioning. Coronal sections spanning the dorsal hippocampus were cut at 40 µm on a cryostat (Leica CM1860) and collected in PBS. Sections were kept in PBS at 4 °C until mounted onto slides and stained.

#### Immunostaining

Free-floating sections were permeabilized for 1 hour at room temperature in 0.3% Triton X-100 with 5% BSA in PBS, then blocked for 2 hours at room temperature in 20% BSA in PBS. Sections were incubated overnight at 4 °C with the primary antibody against NeuN (rabbit anti-NeuN, Abcam Cat. No. AB177487, 1:1000) diluted in blocking buffer, to label neuronal cell bodies. Sections were then washed three times for 15 minutes each in PBS and incubated for 2 hours at room temperature with an Alexa Fluor 647-conjugated secondary antibody raised against rabbit IgG (goat anti-rabbit Alexa Fluor 647, Invitrogen Cat. No. 2433883, 1:1000). Following three additional 15-minutes PBS washes, sections were counterstained with DAPI and cover slipped using Fluoromount-G with DAPI.

#### Confocal Imaging

Stained sections were imaged on a confocal laser scanning microscope (Olympus FV3000, Olympus Corporation) using a 10X objective (UPLSAPO 10X/0.40), in the DAPI (405 nm) and far-red (647 nm, Alexa Fluor 647) channels, operated with Olympus FV31S-SW software. Because a single 10X field of view did not capture the entire hippocampus, tiled images were acquired and stitched into a single composite spanning the ipsilateral and contralateral hippocampus using the FV31S-SW tile-scan feature.

### Quantification of hippocampus histology

Hippocampus histopathology was quantified manually in Fiji (ImageJ2, National Institutes of Health). 10x NeuN images were converted to a maximum intensity projection for analysis. For each animal, 3-6 sections spanning the rostral to dorsal hippocampus were analyzed and averaged. For animals with obliquely cut sections, hemispheres were separated and rostrocaudal levels were manually matched between the IL and CL sides before analysis. All values are expressed as a ratio of IL over CL to account for rostro-caudal differences in hippocampal size and animal-to-animal variability. Hippocampal anatomy was manually quantified by three reviewers. A subset of data was analyzed by all 3 reviewers, demonstrating a high inter-rater reliability with an intraclass correlation coefficient (ICC) of 0.84 and 0.96 for measures of dentate gyrus thickness and CA1 cell layer thickness, respectively.

#### Dentate gyrus granule cell layer (GCL) thickness

Dentate gyrus thickness was measured using the straight-line tool in Fiji, by drawing a line perpendicular to the long axis of the GCL at three points on the upper and three points on the lower blade, which were then averaged to obtain a single thickness value.

#### CA1 pyramidal cell layer (PCL) thickness

As above, CA1 PCL thickness was measured using the straight-line tool in Fiji, and each thickness value represents the average of three separate measurements of the region.

## Results

### KA-induced Status Epilepticus (SE) and Mortality

This study tested several KA injection parameters to optimize the use of synthetic KA in the intrahippocampal kainate mouse model of temporal lobe epilepsy, with the goal of achieving robust induction of status epilepticus with low mortality. The first doses piloted were based on the commonly used doses and injection volumes of 100 or 50nL of 20mM kainic acid^8-11^. Both of these injection parameters were found to have a higher than anticipated mortality (Figure 1), as prior work using natural KA cited acute mortality at <5%^10,12,13^. Acute mortality was calculated here as any death in the first 12 hours post-KA injection, due to either outright death or euthanasia as required by humane endpoints (see Methods). Cumulative 72-hour mortality includes acute mortality and any additional mortality, due to meeting humane endpoints in the following 72-hour period. There was no mortality associated with sham intrahippocampal saline injected controls (n = 10), suggesting that model mortality was specific to KA-induced status epilepticus. The 100nL of 20mM KA injection resulted in an 81.8% acute mortality (n = 11 animals), with no additional cumulative mortality. The 50nL of 20mM KA injection resulted in 15.5% and 31.2% acute and cumulative mortality, respectively (n = 32 animals). Lower concentrations of KA were next tested, including a 50nL injection of 15mM KA (n = 10) and 50nL of 7.5mM KA (n = 2). The 50nL injection of 15mM KA did not reduce mortality compared to 50nL of 20mM injection, with a 10.0% and 30.0% acute and cumulative mortality observed. The 50nL of 7.5mM injection parameters were not explored further due to acute mortality observed in one animal and no induction of status epilepticus observed in a second. These results suggested that titrating based on decreasing concentration of KA alone would not achieve a dose optimal for achieving low mortality while reliably provoking status epilepticus. Conversely, maintaining the total microgram dose at 0.2µg (as in the 50nL of 20mM KA injection) but delivering it in a more dilute formula as 200nL of 5mM KA (n=50) was associated with lower overall mortality (8.0% and 16.0% acute and cumulative mortality) compared to the combined 50nL of 15 or 20mM group (Fisher’s exact test, p = 0.033). Importantly, the 200nL injection group did not differ significantly in duration of status epilepticus (Figure 1B) or cumulative time in Stage 5+ seizures during SE (Figure 1C) compared to the 50nL injection groups.

**Figure 1:**
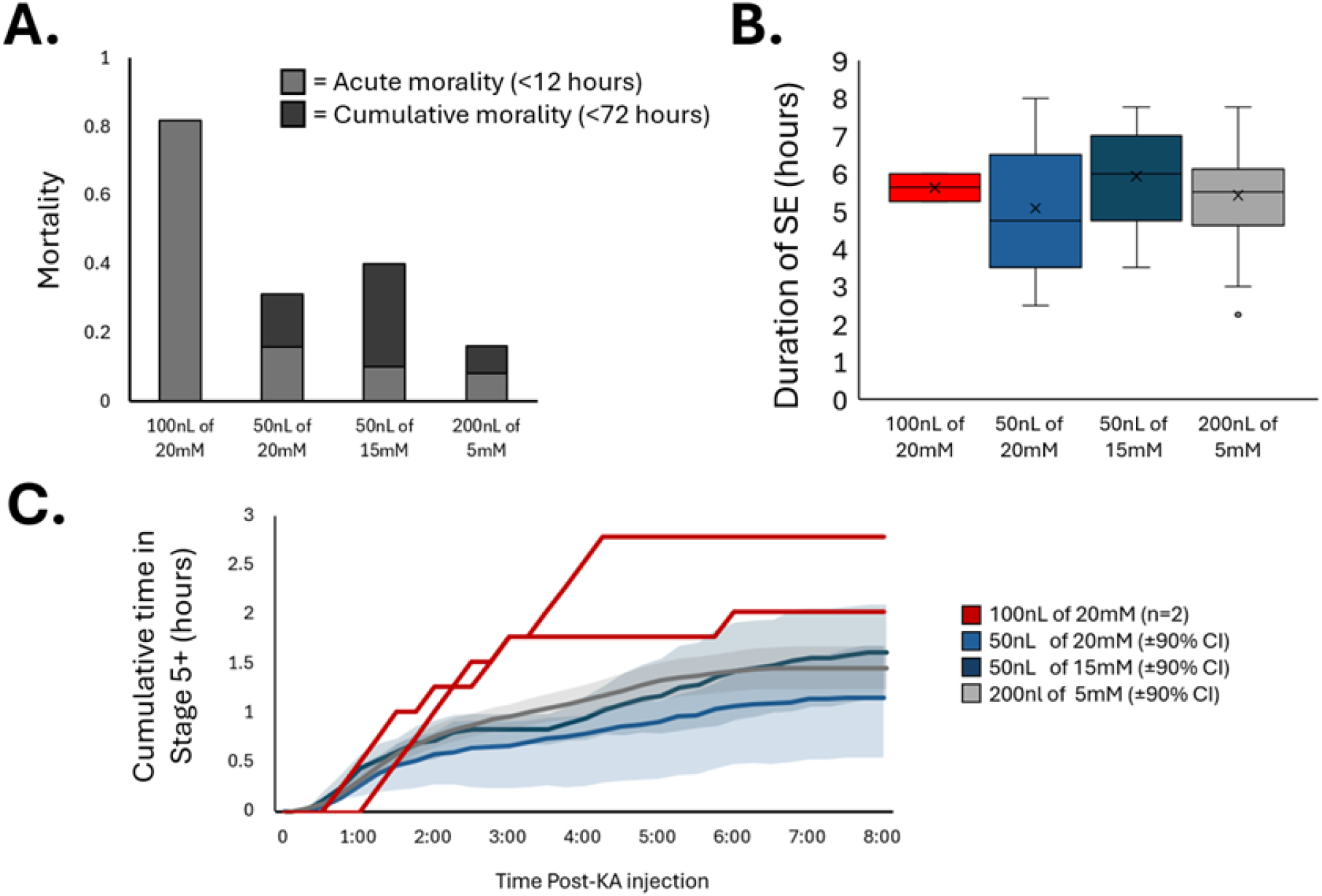
Effect of Kainic Acid Dose and Injection Volume on Mortality and Status Epilepticus. A) Mortality at different injection concentrations and volumes of synthetic kainic acid. Acute mortality included any outright death or euthanasia due to meeting humane endpoints in the first 12 hours after intrahippocampal KA injection. Cumulative morality includes any acute mortality + additional morality in the 72-hour post-KA injection period. (n = 11, 32, 10 and 50 mice, respectively). B) Average duration of KA-induced status epilepticus (SE) as assessed by continuous behavioral monitoring (n = 2, 27, 9 and 49 mice). C) Cumulative time spent in behavioral seizure of Stage 5 or greater post-KA injection. (n = 2 individual animals at 100nL of 20mM KA dose). Remaining lines are mean +/-90% confidence intervals (n = 27, 9 and 49, respectively). Seizure staging was performed in 15 minutes increments by behavioral screening, where each 15-minute epoch was assigned a seizure severity score (on modified Racine scale of 0-6) based on the most severe seizure observed in that period.

### IHK Histopathology

Temporal lobe epilepsy is associated with hippocampal sclerosis, with common features of CA1 cell loss and granule cell layer dispersion in the dentate gyrus^21,22^. Here we quantified these anatomical features from NeuN immunostained sections from sham IH-saline injected controls (n = 4) and IHK animals receiving 50nL of 15 or 20mM KA (n = 11), or 200nL of 5mM KA (n = 8). Tissue was collected for immunostaining at least 2-weeks post KA-injection (on average, 24.6 ± 2.3 days post KA-injection). CA1 cell loss was quantified by measuring the thickness of the CA1 pyramidal cell layer and calculating the ratio of ipsilateral to contralateral thickness. 3-6 sections were analyzed per animal and averaged to obtain a single CA1 thickness ratio for each animal. CA1 cell layer thickness ratio was found to be 0.95 ± 0.08, 0.43 ± 0.08, and 0.69 ± 0.07, for sham, 50nL and 200nL groups, respectively (Figure 2). KA-injection resulted in significant ipsilateral CA1 cell loss compared to sham-injected controls (Welch’s t test, t = 4.12(6.9), p = 0.004). The 50nL group was associated with greater CA1 cell loss compared to the 200nL group (Welch’s t test, t = -2.44(17.0), p = 0.03), suggesting that 50nL of 20/15mM KA injection was more injurious. Granule cell layer dispersion was quantified by taking the ratio of the ipsilateral to contralateral granule cell layer thickness, with 3-6 sections per animal, as above. KA-injection was associated with a significant increase in the thickness of the granule cell layer consistent with KA-induced granule cell layer dispersion (Welch’s test, t = 3.98(20.3), p < 0.001), without a significant difference between the 50nL and 200nL KA groups. These results demonstrate that either the 50nL or 200nL injection protocol results in stereotypical TLE-like hippocampal histopathology, with less overall CA1 cell loss observed in the 200nL injection cohort.

**Figure 2:**
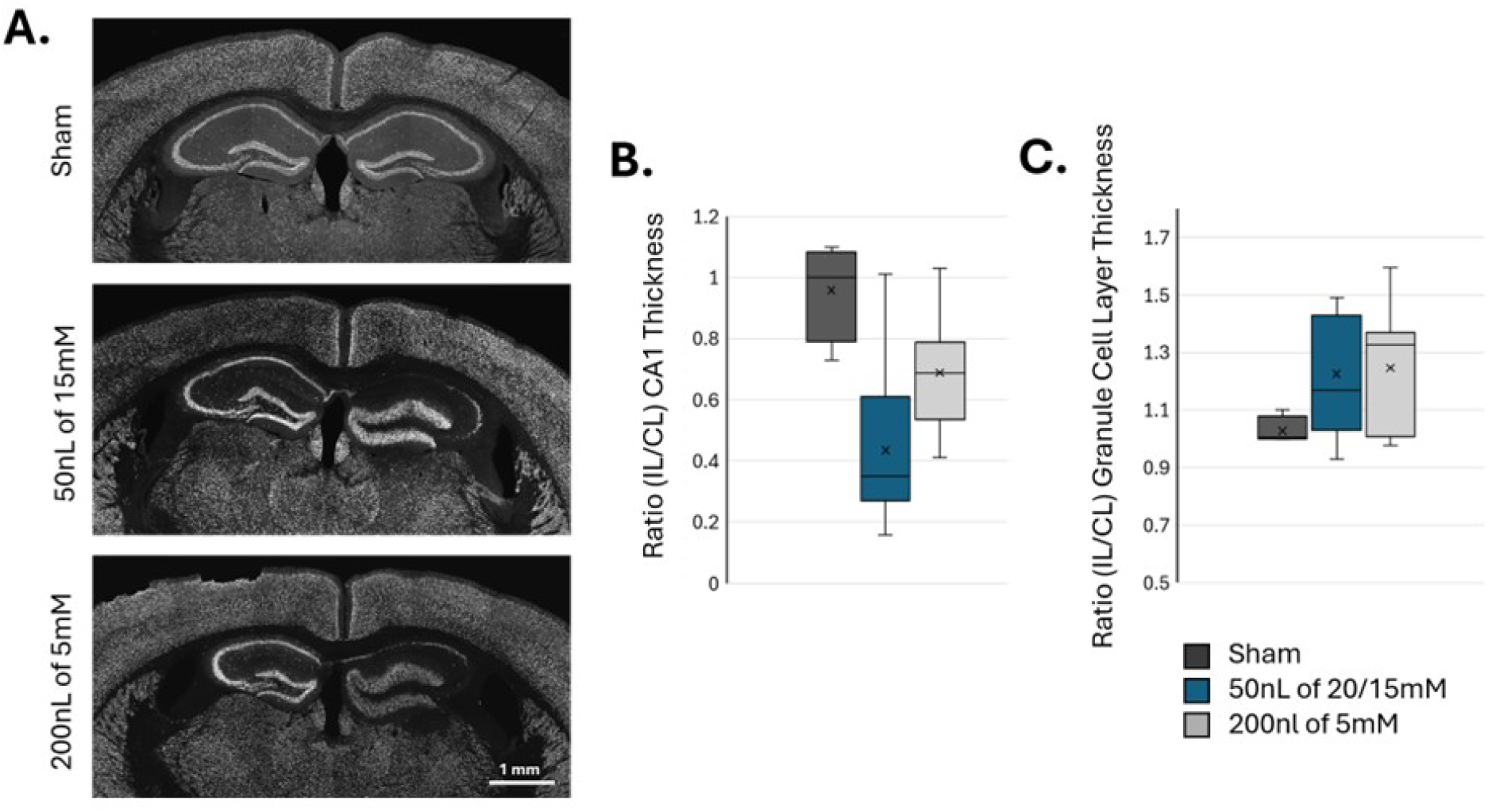
TLE-Associated Hippocampal Histopathology Across Kainic Acid Injection Volumes. A) Representative images of NeuN immunostaining of sham (intrahippocampal saline injected), IHK with 50nL of 15mM KA and IHK with 200nL of 5mM KA animal. B) Boxplot of CA1 cell layer thickness, expressed as ratio of ipsilateral (KA-injected) over contralateral. (n = 4 saline-injected sham animals, 11 animals receiving 50nL of 15 or 20mM KA and 8 animals receiving 200nL of 5mM KA). KA-injection resulted in significant CA1 cell layer thinning (p < 0.01, Welch’s t test, KA groups vs sham). 50nL group was associated with greater CA1 cell loss compared to 200nL group (p=0.03, Welch’s t test). C) Boxplot of ratio of dentate gyrus granule cell layer thickness, expressed as ratio of ipsilateral over contralateral. KA-injection was associated with significant granule cell layer dispersion (p < 0.001, Welch’s t test, KA groups vs sham), without a significant difference between the 50nL and 200nL injection groups.

### IHK spontaneous seizures

In order to assess the frequency of spontaneous seizures, a cohort of 50nL of 20mM KA injected and 200nL of 5mM injected animals were implanted with cortical EEG and underwent at least one-week of chronic video-EEG recording. Both KA injection groups displayed robust development of epileptiform and spontaneous seizure activity. Seizures were identified by high-amplitude EEG spiking accompanied by progressive evolution in waveform amplitude and morphology, lasting >10 seconds, typically followed by post-ictal depression (Figure 3). 97% of observed seizures were associated with convulsive behaviors, with severity scores of 4-6 on the modified Racine scale (Table 1)^18,19^. The remaining electrographic seizures were similar in duration and EEG appearance but were not accompanied by correlated convulsive behavior. The mean seizure frequency was 1.33 seizures/day ± 0.89 and 2.89 seizures/day ± 1.02, in the 50 nL (n = 6) and 200nL injection groups (n = 7), respectively. (One IHK animal in the 50nL cohort did not have a spontaneous seizure during the one-week video-EEG recording period, but did present with multiple, high-amplitude, rhythmic spiking events. Given the relatively short recording duration, the absence of observed seizures in this animal does not exclude the possibility of epilepsy^23^.) Although the study was not powered to detect potentially modest differences in seizure frequency between KA injection protocols, these results demonstrate that both KA-injected groups reliably produce an epileptic phenotype. Notably, seizure frequency varied substantially between individual animals within both injection groups. Animals also differed markedly in the temporal distribution of seizures, with some exhibiting pronounced clustering of events and others showing relatively regular seizure occurrence with little evidence of clustering (Figure 4A). The variance-to-mean ratio (VMR) of daily seizure counts was similar between injection groups, at 1.55 ± 0.68 and 1.39 ± 0.39 for the 50nL and 200nL groups, respectively (Figure 4B). Together, these findings highlight substantial animal-to-animal variability in both seizure frequency and temporal organization, underscoring the need for caution when interpreting pooled seizure measures across animals or experimental conditions.

**Figure 3:**
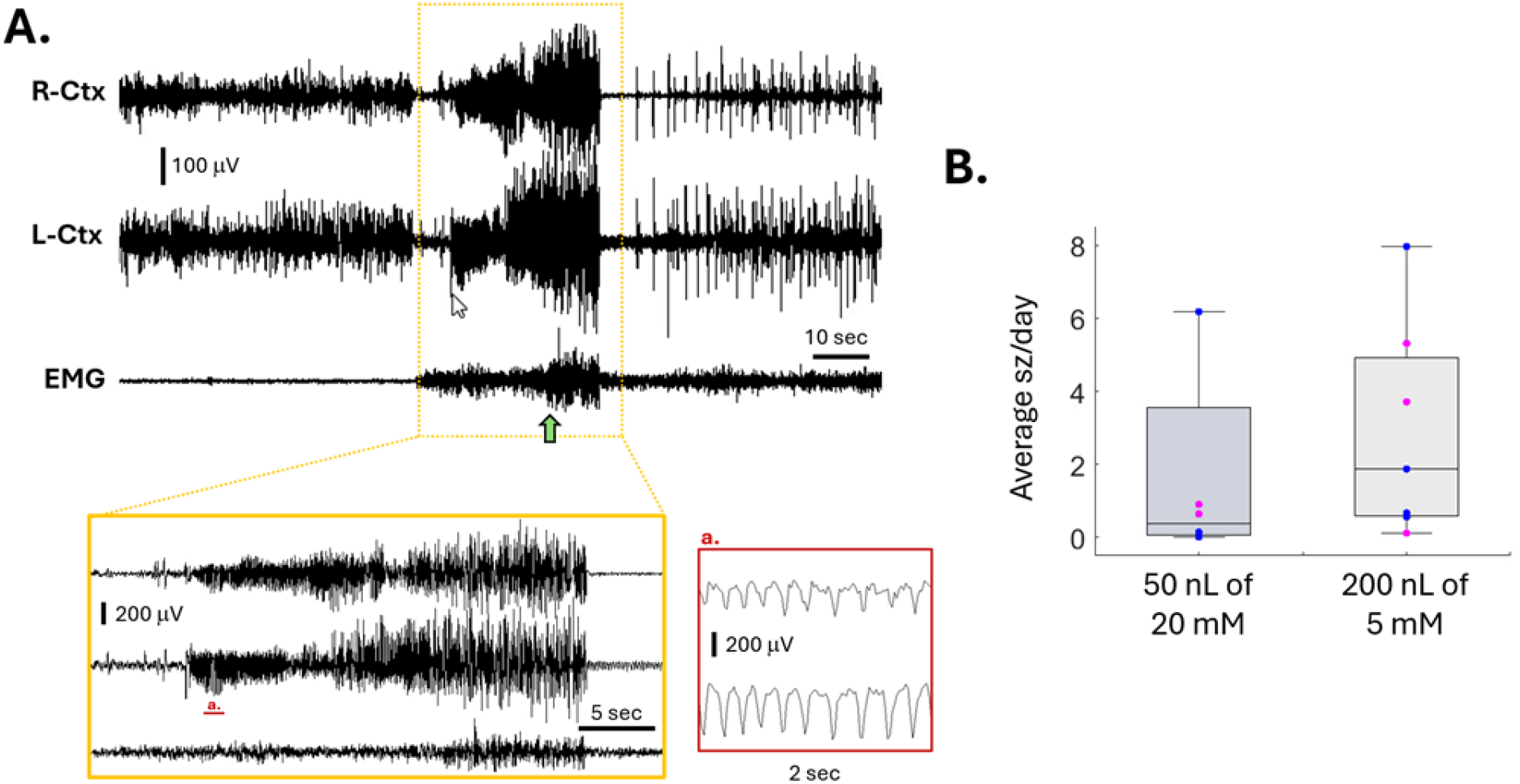
Robust Induction of Spontaneous Recurrent Seizures Across Kainic Acid Injection Volumes. A) Representative EEG recording of spontaneous convulsive seizure in an IHK animal receiving a 200nL injection of 5mM KA to the right hippocampus, with electrodes placed in the right cortical area (R-Ctx) and left cortical area (L-Ctx), and EMG recording from nuchal muscles. Green arrow indicates time that behavioral convulsions started. B) Boxplot of mean seizures per day, as quantified from 7-9 days of continuous video-EEG per animal, for 50nL of 20mM KA group (n = 6) and 200nL of 5mM KA group (n=7). Blue dots indicated values from male mice and magenta dots indicate values from female mice.

**Figure 4:**
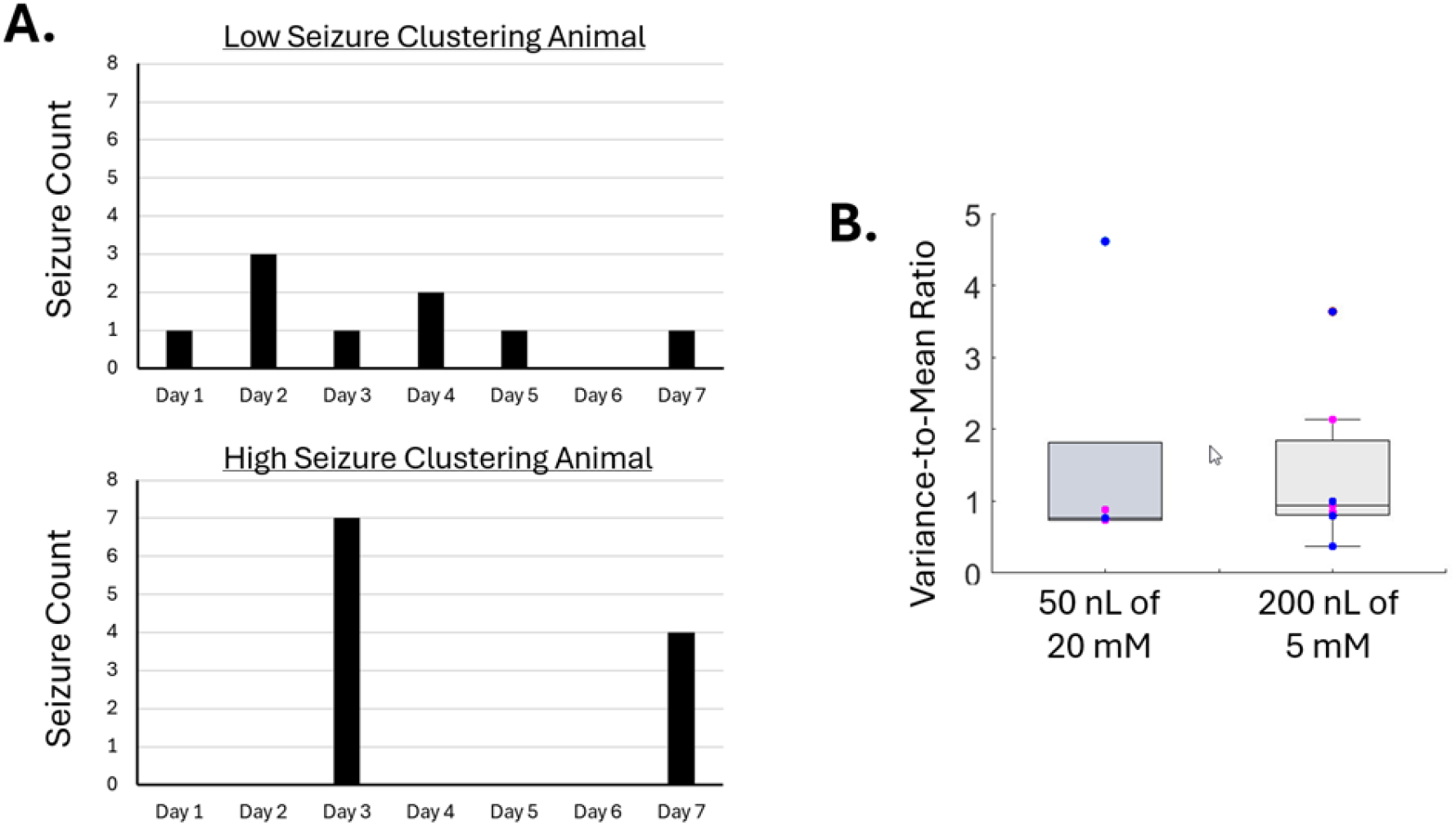
Seizure Clustering Across Kainic Acid Injection Volumes. A) Representative examples of per day seizure counts for IHK animals with low seizure clustering (top) and high seizure clustering (bottom). B) Boxplot of variance-to-mean ration, as quantified from daily seizure counts over 7-9 days of continuous video-EEG per animal, for 50nL of 20mM KA group (n = 6) and 200nL of 5mM KA group (n=7). Blue dots indicated values from male mice and magenta dots indicate values from female mice.

## Discussion

In conclusion, intrahippocampal injection of synthetic KA delivered at a variety of concentrations and volumes was found to reliably induced status epilepticus. Interestingly, while mortality could be titrated by lowering the concentration and total amount of KA administered, the duration and severity of status epilepticus was relatively insensitive to these changes. Notably, comparison of injection protocols with similar total amounts of KA suggests that distributing the KA over a wider volume may be more important for reducing mortality than reducing the total KA amount alone. Although both concentrations used here (20mM and 5mM) are substantially above the reported EC50 for KA receptors^24^, the higher-concentration injection potentially produces a higher peak local concentration and steeper concentration gradient near the injection site. A larger injection volume may therefore distribute KA more broadly and limit the severity of the focal exposure. Additionally, because the present experiments used a synthetic KA preparation, whereas many earlier works used naturally derived KA, differences in formulation or preparation could also contribute to differences between our work and previous studies. However, regardless of the basis for these differences, we propose that the 200nL of 5mM synthetic KA injection protocol effectively balances low mortality, with robust induction of status epilepticus and development of hippocampal sclerosis and recurrent spontaneous seizures.

Interestingly, we observed a high degree of heterogeneity in the seizure profile of individual animals. In addition to KA injection parameters, as considered here, variability in epileptic phenotypes may be influences by additional factors such as sex, genetic background and stage of disease progression. In the current study, our relatively small cohort size precluded comparisons of these factors but it would be of interest to perform larger, factorially designed studies to determine whether these biological variables contribute to the heterogeneity observed here. Although inter-animal variability represents a challenge for experimental design and statistical power, it may also provide an opportunity to investigate how biological differences give rise to distinct seizure phenotypes and potentially capture aspects of the clinical heterogeneity of epilepsy. Additionally, the pronounced seizure clustering observed in this study underscores the importance of sufficiently long recording periods to obtain reliable estimates of seizure burden and should be considered when designing and interpreting experiments examining manipulations of seizure frequency. More generally, the high level of variation in seizure dynamics associated with the IHK model needs to be accounted for when comparing seizure profiles and outcomes across studies.

A further consideration when comparing seizure frequency across studies is the definition used to identify seizure events. Some studies of the IHK model have classified brief episodes of electrographic spiking, sometimes lasting only a few seconds, as seizures^10,11,25-27^. In contrast, the present study defined seizures as sustained high-amplitude events with electrographic evolution lasting >10 s, in line with the seizure definitions used in other labs^8,28^. The high frequency (>20/hr) of the brief epileptiform events observed in this model does offer advantages for studies in which seizure-like activity is used as a sensitive readout of experimental manipulations. However, these events should not be considered interchangeable with sustained electrographic or convulsive seizures, as they likely represent distinct forms of pathological network activity.

Finally, several limitations of the present study should be considered. First, the relatively small cohort sizes limited our ability to determine the contributions of biological factors such as sex, genetic background, and disease progression to the observed variability (although, these factors were balanced between the two injection groups in this study to minimize potential confounding effects). Second, the duration of EEG monitoring was relatively short and therefore may not fully capture the temporal organization of seizures, particularly in animals with pronounced seizure clustering. Longitudinal studies with extended recording periods and repeated assessment at different stages of disease progression will be important to determine the stability of seizure phenotypes over time. More broadly, these considerations highlight the importance of carefully controlling and reporting experimental parameters that may contribute to variability in the IHK model and animal model of epilepsy in general. Careful definition and reporting of model conditions will facilitate reproducibility and meaningful comparison across studies. Overall, by establishing conditions for the use of synthetic KA that robustly recapitulate key features of TLE, this work strengthens the utility of the IHK model as a platform for investigating the mechanisms underlying epilepsy and its diverse seizure phenotypes.

## Acknowledgements

This work was supported by National Institutes of Health (R01NS112538 to K.P.L, and R35NS116852 to K.J.S.) and CURE Epilepsy (Taking Flight Award to L.A.L.).

